# Module analysis captures pancancer genetically and epigenetically deregulated cancer driver genes for smoking and antiviral response

**DOI:** 10.1101/216754

**Authors:** Magali Champion, Kevin Brennan, Tom Croonenborghs, Andrew J. Gentles, Nathalie Pochet, Olivier Gevaert

## Abstract

The availability of increasing volumes of multi-omics profiles across many cancers promises to improve our understanding of the regulatory mechanisms underlying cancer. The main challenge is to integrate these multiple levels of omics profiles and especially to analyze them across many cancers. Here we present AMARETTO, an algorithm that addresses both challenges in three steps. First, AMARETTO identifies potential cancer driver genes through integration of copy number, DNA methylation and gene expression data. Then AMARETTO connects these driver genes with co-expressed target genes that they control, defined as regulatory modules. Thirdly, we connect AMARETTO modules identified from different cancer sites into a pancancer network to identify cancer driver genes. Here we applied AMARETTO in a pancancer study comprising eleven cancer sites and confirmed that AMARETTO captures hallmarks of cancer. We also demonstrated that AMARETTO enables the identification of novel pancancer driver genes. In particular, our analysis led to the identification of pancancer driver genes of smoking-induced cancers and ‘antiviral’ interferon-modulated innate immune response.

**Software availability:** AMARETTO is available as an R package at https://bitbucket.org/gevaertlab/pancanceramaretto

**Highlights:**

- We present an algorithm for pancancer identification of cancer driver genes based on multiomics data fusion
- GPX2 is a novel driver gene in smoking induced cancers and validated using knockdown of GPX2 in the A549 cell line.
- OAS2 is a novel driver gene defining cancers with an antiviral signature supported by increased infiltration of tumor-associated macrophages.

**Research in context:** We present an algorithm that combines multiple sources of molecular data to identify novel genes that are involved in cancer development. We applied this algorithm on multiple cancers in a combined fashion and identified a network of pancancer driver genes. We highlighted two genes in detail GPX2 and OAS2. We showed that GPX2 is an important cancer gene in smoking induced cancers, and validated our predictions using experimental data where GPX2 was inactivated in a lung cancer cell line. Similarly we showed that OAS2 is an important cancer driver gene in cancers that show an antiviral signature.

## Introduction

In the last two decades, advances in high-throughput experimental technologies have produced an abundance of molecular data. An increasing number of large multi-omics projects have launched and provide millions of data points for thousands of biological samples. For example, The Cancer Genome Atlas (TCGA) project ^1-3^ was launched to improve our ability to diagnose, treat and prevent cancer and has produced an enormous amount of multi-omics data. Interpreting these high dimensional datasets to identify novel cancer driver genes represents an outstanding challenge. True cancer driver genes are those whose perturbation pushes a cell towards a malignant phenotype. Within this study, we define cancer driver genes as genes that fulfill all of the following criteria: (1) genes that are genetically and/or epigenetically deregulated in cancer, (2) genes whose genetic and epigenetic aberrations have a direct impact on their own functional gene expression levels, and (3) genes that are predicted to play regulatory roles high in the causal hierarchy of the origin of tumors. These include, for example, transcription factors, cell cycle genes or epigenetic modifying enzymes, whose altered state in cancer results in deregulation of downstream target genes; as well as upstream signaling molecules. They typically hide amongst a large number of passenger genes that are only by chance genetically or epigenetically altered ^4^.

Previously, several computational methods have been developed to integrate multi-omics data. For example, Ciriello et al. used a method based on mutual exclusivity of copy number and mutation events to identify driver genes in glioblastoma ^5^. Similarly, Vandin et al. developed a method to identify driver genes in cancer, but focused on finding pathways with a significant enrichment of mutually exclusive genes ^6^. In addition, Akavia et al. built further on this work and used copy number data to identify potential cancer driver genes in a modified Bayesian module network analysis called CONEXIC ^7^. More recently, other groups are focusing on identifying driver genes through network analysis of copy number data to identify potential drivers using a Bayesian module network analysis ^8^. We have previously developed AMARETTO, an algorithm that integrates copy number, DNA methylation and gene expression data to identify a set of driver genes altered by DNA methylation or DNA copy number alterations, and constructs a gene expression network to connect them to clusters of co-expressed genes, defined as modules ^9, 10^. These gene expression modules are subsequently ascribed biological pathways using gene set enrichment analysis (GSEA), revealing the pathways affected by cancer driver gene regulation. AMARETTO is thus a data driven pathway approach, using genomic, epigenomics and transcriptomics data as inputs, and produces modules and cancer driver genes associated with these modules as output. Integration of epigenomics data is essential to comprehensive analysis of cancer genomic analysis, as DNA methylation is a major mechanism of transcriptional deregulation in virtually all cancers. For example, cancer driver genes such as BRCA1 and MLH1, which are often altered by mutation in cancer, are also frequently deregulated by DNA methylation in other patients, with similar downstream consequences ^11-13^. Our data-driven pathway approach contrasts with previous work that relies upon use of known cancer pathways and networks such as PARADIGM, an algorithm that uses human-curated pathways and estimates their activity using DNA copy number and mRNA expression data ^14^.

Here, we present an extension of AMARETTO to a pancancer application using multi-omics data of eleven cancer sites from TCGA. We show that AMARETTO captures modules enriched in major pathways of cancers and modules that accurately predict molecular subtypes. Next, we connect the modules of co-expressed genes in a pancancer module network. We show that this allows the identification of major oncogenic pathways and cancer driver genes involved in multiple cancers. More specifically, we identified a pancancer driver gene that is involved in smoking induced cancers and a pancancer driver gene that is involved in antiviral IFN modulated immune response. Overall, our results show the potential of pancancer multi-omics data fusion to identify cancer drivers that are high within the causal hierarchy of cancer development and associated with common pathways across different types of tumors that eventually can lead to the identification of pancancer drug targets. The AMARETTO algorithm and its pancancer application are publicly available.

## Materials and Methods

### Data preprocessing

We used gene expression, copy number and DNA methylation data from TCGA for 11 cancer sites, namely bladder urothelial carcinoma (BLCA), breast invasive carcinoma (BRCA), colon and rectal adenocarcinoma (COADREAD), glioblastoma (GBM), head and neck squamous cell carcinoma (HNSC), clear cell renal carcinoma (KIRC), acute myeloid leukemia (LAML), lung adenocarcinoma (LUAD), lung squamous cell carcinoma (LUSC), ovarian serous cystadenocarcinoma (OV) and uterine corpus endometrial carcinoma (UCEC) (Table 1). All data sets are available at the TCGA data portals.

**Table 1.**

| TCGA cancer site | TCGA cancer code | Gene Expression |  | GISTIC |  | MethylMix |  |
| --- | --- | --- | --- | --- | --- | --- | --- |
|  |  | Samples | Genes | Samples | Genes<sup>1</sup> | Samples | Genes<sup>2</sup> |
| Bladder urothelial carcinoma | BLCA | 181 | 15.432 | 178 | 1.974 | 123 | 472 |
| Breast invasive carcinoma | BRCA | 985 | 16.02 | 968 | 1.523 | 887 | 890 |
| Colon and rectum adenocarcinoma | COADREAD | 589 | 15.533 | 578 | 2.523 | 570 | 522 |
| Glioblastoma multiforme | GBM | 501 | 17.811 | 481 | 1.561 | 321 | 395 |
| Head and neck squamous cell carcinoma | HNSC | 371 | 15.828 | 365 | 2.184 | 308 | 753 |
| Kidney renal clear cell carcinoma | KIRC | 509 | 16.123 | 501 | 3.052 | 497 | 567 |
| Acute myeloid leukemia | LAML | 173 | 14.296 | 166 | 1.681 | 170 | 613 |
| Lung adenocarcinoma | LUAD | 489 | 16.092 | 487 | 3.585 | 367 | 678 |
| Lung squamous cell carcinoma | LUSC | 490 | 16.219 | 487 | 2.592 | 355 | 679 |
| Ovarian serous cystadenocarcinoma | OV | 541 | 17.814 | 528 | 1.499 | 540 | 510 |
| Uterine corpus endometrial carcinoma | UCEC | 508 | 15.706 | 500 | 2.074 | 496 | 821 |

<sup>1</sup> Number of significant genes found after running GISTIC (data available in the TCGA data portal).
<sup>2</sup> Number of genes with significant methylation patterns identified using MethylMix.

The gene expression data were produced using Agilent microarrays for GBM and OV cancers, and RNA sequencing for all other cancer sites. Preprocessing was done by log-transformation and quantile normalization of the arrays. The DNA methylation data were generated using the Illumina Infinium Human Methylation 27 Bead Chip. DNA methylation was quantified using β-values ranging from 0 to 1 according to the DNA methylation levels. We removed CpG sites with more than 10% of missing values in all samples. We used the 15-K nearest neighbor algorithm to estimate the remaining missing values in the data set ^15^. Finally, the copy number data we used are produced by the Agilent Sure Print G3 Human CGH Microarray Kit 1Mx1M platform. This platform has high redundancy at the gene level, but we observed high correlation between probes matching the same gene. Therefore, probes matching the same gene were merged by taking the average. For all data sources, gene annotation was translated to official gene symbols based on the HUGO Gene Nomenclature Committee (version August 2012). TCGA samples are analyzed in batches and significant batch effects were observed based on a one-way analysis of variance in most data modes. We applied Combat to adjust for these effects ^16^.

### AMARETTO: multi-omics data fusion

Our approach for analyzing TCGA cancer data is based on AMARETTO, a novel algorithm devoted to construct modules of co-expressed genes through the integration of multi-omics data ^9, 10^. More precisely, AMARETTO is a three-step algorithm that (i) identifies tumor specific DNA copy number or DNA methylation changes, (ii) identifies a set of potential cancer driver genes by integrating DNA copy number, DNA methylation and gene expression data, (iii) connects these cancer driver genes to modules of co-expressed target genes that they control using a penalized regulatory program. AMARETTO, consists of three steps (Figure 1):

**Figure 1:**
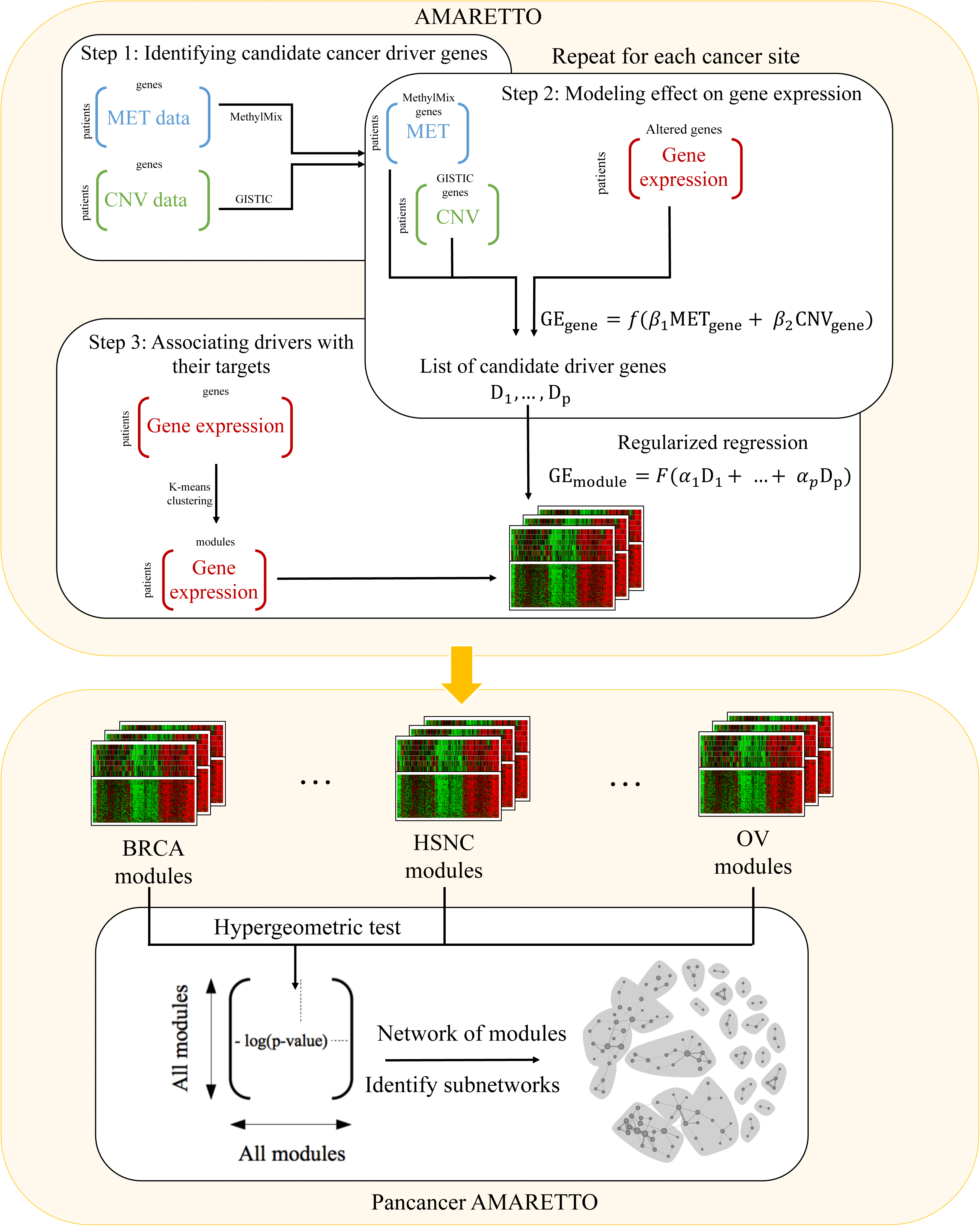
Workflow of pancancer AMARETTO analysis. AMARETTO generates a list of candidate cancer driver genes by investigating significant correlations between methylation, copy number and gene expression data for each putative driver gene separately, and connects them with their downstream targets by constructing a module network. We then identify modules that capture major hallmarks of cancers. We finally detect subnetworks of modules across all cancers.

Step 1: Identification of candidate cancer driver genes with tumor-specific DNA copy number or DNA methylation alterations compared to normal tissue: we first restrict the list of candidates to genes that have either copy number or DNA methylation alterations. These alterations are detected using the GISTIC ^17, 18^ and MethylMix ^19, 20^ algorithms for copy number and DNA methylation data respectively. GISTIC separately models arm-level and focal alterations, identifying amplified and deleted genes. Modeling DNA methylation aberrations in cancer is less well studied. We recently developed MethylMix, a method that identifies hypo and hypermethylated genes by (i) detecting methylation states of each gene with univariate beta mixture models, (ii) comparing them with the DNA methylation levels of normal tissue samples. We used GISTIC to identify significantly and recurrently deleted or amplified regions in the genome ^18^. Similarly, we used MethylMix to identify recurrently hyper-or hypomethylated genes ^19^.

Step 2: Modeling effect of candidate driver genes on gene expression: a given gene is considered as a candidate driver gene if its expression can be explained by genomic events. Our rationale is that genes driven by multiple genomic events in a significant subset of samples are unlikely to be randomly deregulated. To establish a list of cancer driver genes, we thus investigate the linear effects of copy number and DNA methylation on gene expression through a linear regression model performed on each gene independently:
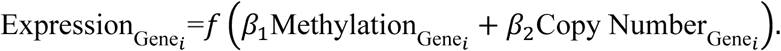

We used the R^2^ statistic to evaluate whether copy number has a significant positive effect (*β*_2_ > 0) and DNA methylation a significant negative effect (*β*_1_ < 0) on gene expression.

Step 3: Associating candidate driver genes with their downstream targets: given the cancer driver genes identified in Step 1, Step 2 aims at connecting them to their regulated targets to construct the regulatory module network. First, the filtered data are clustered in modules of co-expressed genes using a k-means algorithm with 100 clusters. Then, we learn the regulatory programs for each of the modules as a linear combination of cancer driver genes that together explain each module’s mean expression:
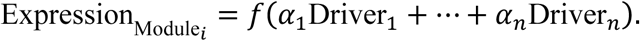

In order to induce sparseness, we use an L_!_-penalty on the regression weights ^21^. The modules are further optimized by running iteratively over the two following steps: (i) reassigning genes based on the closest match to the updated regulatory programs, (ii) updating the regulatory programs based on the new gene assignments ^22^. These two steps are repeated until less than 1% of the cancer driver genes are assigned to new modules.

### Pancancer module network

After running AMARETTO on each cancer site individually, the modules are connected in a pancancer network. Specifically, we evaluate whether there is a significant association between all pairs of modules through a hyper-geometric test. We correct for multiple hypothesis testing using the false discovery rate ^23^. We consider the association to be significant if both of the following conditions are satisfied: (i) the adjusted p-value is smaller than 0.05 and (ii) the overlap between two modules is larger than 5 genes. This defines a module network where each edge is scored based on the negative log-transformed adjusted p-value.

We cluster the weighted module network to identify significantly connected subnetworks using the Girvan-Newman algorithm ^24^. This algorithm is a divisive method, which aims at detecting subnetworks by progressively removing edges from the module network according to a score. The original proposed score is based on the betweenness of edges, where betweenness is a measure that favors edges that are between subnetworks, and thus responsible for connecting many pairs of others. We used the igraph R package to visualize the network and the edge.betweenness.community function (implementation of the Girvan-Newman algorithm) to detect subnetworks. We only focus on subnetworks that satisfy the following conditions: number of nodes larger than the 1% of the total number of nodes in the network, number of represented cancers larger than the 10% of the subnetwork size (and at least, larger than 2) and finally, ratio between edges inside/outside the subnetwork larger than ½.

### Gene set enrichment analysis

To assign biological meaning to these subnetworks of modules, we perform gene set enrichment analysis based on the databases GeneSetDB ^25^ and MSigDB ^26^. For the latter, we restrict the enrichment to hallmark (H), curated (C2), GO (C5), oncogenic (C6) and immunologic signatures (C7) gene sets, which include the gene sets most relevant to cancer gene expression profiles. The enrichment is evaluated by performing a hyper-geometric test, corrected for multiple testing using the FDR ^23^. We used averaged p-values to combine p-values of the pathway enrichment for modules within a subnetwork. We used the following cutoffs: p-value smaller than 0.05, overlap with one module larger than 5 and gene set size smaller than 500. As a negative control we compared the enrichment of the actual modules for each cancer with 100 random permutations of the gene to module assignments. We then counted for each permutation the average number of significant gene sets per module, over all modules using the same cutoffs as the actual enrichment results.

### Prediction performances

After running AMARETTO on the provided data sets, we computed the prediction performances by comparing the observed module expression data matrix with its predicted value. We reported the averaged mean squared error (MSE) and the R-square taken across all modules.

### Smoking signatures

To investigate the role played by GPX2 on smoking related pathways of different cancer sites, we first used an oxidative gene signature from the Gene Ontology (Supplementary Table 6a). We defined an associated score by taking the average expression of these genes. We then used a Pearson test to measure the correlation with GPX2 expression. We did the same for a second GO signature associated to xenobiotic metabolism (Supplementary Table 6a).

### Correlation with smoking data

We investigated whether GPX2 expression is correlated with smoking. We used clinical data from TCGA containing 743 characteristics (e.g. ethnicity, gender, tumor size…). We restrict our study to clinical variables that are related to smoking (profile, started smoking year, stopping smoking year and pack years). We obtained a significant number of clinical data for only 4 of the 11 cancer sites, namely BLCA, HNSC, LUAD and LUSC. For each of these cancer sites, correlation coefficients between the associated clinical variables and GPX2 expression are calculated through the Spearman test for continuous variables, and the Wilcoxon test, or Kendall test for discrete variables with two or more than two groups respectively. In addition, we drew boxplots representing the association between smoking profile and GPX2 expression.

### Experimental validation using GPX2 knockdown experiments

We extracted GPX2 perturbation experiments from the LINCS database ^27, 28^. GPX2 perturbation experiments were available for the lung adenocarcinoma A549 cell line that best resembles the LUAD cancer site, while no matching cell lines were available for the 4 other cancer sites, i.e., LUSC, BLCA, HNSC and UCEC. Four perturbation experiments measuring experimental targets of GPX2 knockdown in the A549 cell line were available in LINCS, including three shRNA experiments and a consensus signature derived from these three experiments. In one of these four experiments, a positive differential expression z-score for GPX2 was measured, and we therefore removed this experiment from the analysis since successful GPX2 knockdown is expected to result in a negative z-score. We used the “preranked” Gene Set Enrichment Analysis (GSEA) (GSEA) ^29^ tool to test for enrichment of the target genes of modules regulated by GPX2 in the genome-wide differential expression z-score profiles of the three GPX2 knockdown experiments. We restricted our analysis to the landmark genes (measured on the L1000 platform) and bing genes (inferred with high confidence), and we collapsed multiple probes to unique genes by selecting the probe with the most reliable (landmark over bing) and the highest absolute z-score value. GSEA enrichments were estimated using the normalized enrichment score (NES) and significance of the enrichments was assessed at the FDR<0.25 level as well as p-value<0.05 and FDR<0.25 levels.

### Correlation with PDL1-PDL2 expression

To investigate the role played by OAS2 on immune response pathways of different cancer sites, we correlated OAS2 expression with CD274, more commonly known as PDL1, and PDCD1LG2 expressions, more commonly known as PDL2, using a Pearson test.

### Inference of tumor associated leukocyte levels using CIBERSORT

CIBERSORT ^30, 31^ is a computational method that characterizes cell composition of complex tissues from their expression profiles. We applied CIBERSORT to TCGA gene expression data to infer leukocyte representation in the 11 considered cancer sites. More precisely, we used expression profiles for 22 distinct leukocyte subsets (TALs). Only patients for whom estimated p-values are less than 0.05 (indicating high confidence TAL estimation) were included in downstream analyses.

## Results

AMARETTO is an algorithm that allows the integration of multi-omics data to identify cancer drivers and associate them to their downstream targets. Here, we present a pancancer application of AMARETTO on eleven different cancer sites (Figure 1, Table 1).

### AMARETTO captures major hallmarks of cancer

AMARETTO modules capture the major oncogenic pathways whereas randomly permuted modules do not result in significant gene sets in all cancer sites (Figure 2, Supplementary Table 1). We found 22 modules enriched in cell cycle pathways. Four of these modules are regulated by CHEK1, a well-known cell cycle gene required for checkpoint-mediated cell cycle arrest in response to DNA damage ^32, 33^. Next, we found that 43 modules are highly enriched with genes related to angiogenesis. The most common cancer driver gene, FSTL1, regulates 8 of the 43 enriched modules, representing a potential cancer driver gene that regulates angiogenesis. This gene has been shown to be involved in proliferation, migration and invasion ^34-36^ and was recently linked with angiogenesis in post-myocardial infarction rats ^37^. Twelve modules are enriched in epithelial-to-mesenchymal transition (EMT) pathways. The cancer driver genes of EMT modules include TGFB3, a member of the TGF beta pathway known to regulate EMT ^38-41^, and NUAK, which has been implicated in several cancers ^42-44^. In addition, NUAK1 is involved in EMT in ovarian cancer ^45^ and is part of two ovarian cancer modules identified by AMARETTO that are enriched in EMT-related genes ^38^. Next, 100 modules are enriched in immune response pathways, with the inflammatory chemokine CCL5 regulating 14 of these modules ^46^. Overall, AMARETTO enabled us to find modules enriched in many major hallmarks of cancers, including hypoxia, apoptosis, metastases, integrin and epidermal growth factor receptor (EGFR) signaling demonstrating the validity of our approach (Figure 2, Supplementary Table 1).

**Figure 2:**
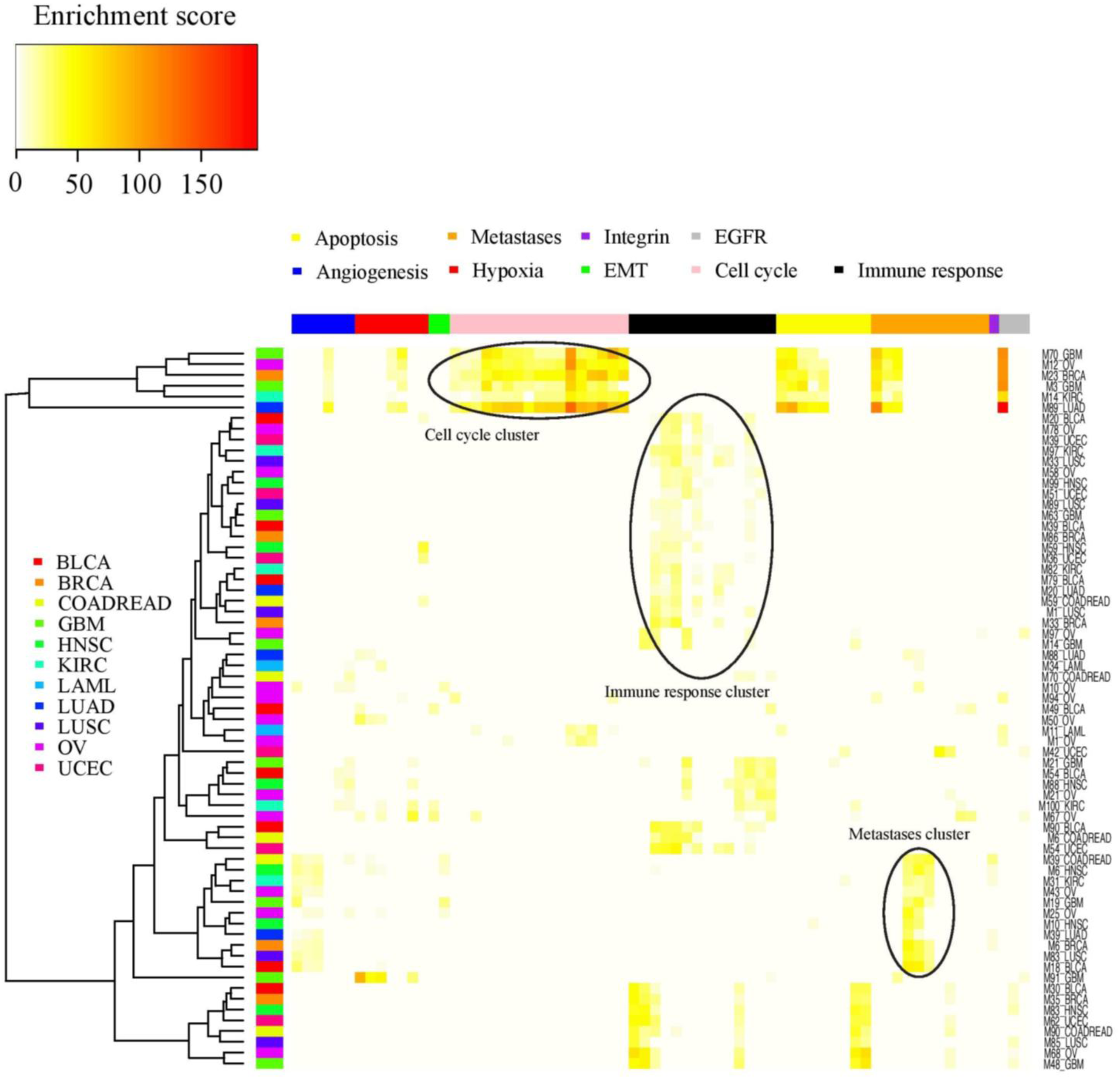
Heatmap representing the enrichment between modules of all cancer sites (in rows) and gene sets associated with major hallmarks of cancers (in columns): angiogenesis, hypoxia, epithelial mesenchymal transition (EMT), cell cycle, immune response, apoptosis, metastases, integrin and epidermal growth factor receptor (EGFR).

### Methylation driven genes are more predictive of downstream expression than DNA copy number driver genes

The module networks for each of the 11 cancer types contain on average 408 cancer driver genes and 7.67 cancer driver genes per module. The top cancer driver genes across all histologies include 45 genes that regulate more than 15 modules across an average number of 4.9 cancer sites per gene (Supplementary Table 2). Interestingly, for all cancers, a higher proportion of selected drivers are DNA methylation driven compared to DNA copy number driven genes (Supplementary Figure 1). Over 90% of cancer driver genes present aberrant DNA methylation patterns, highlighting the importance of DNA methylation-mediated deregulation. Moreover, using methylation data with or without copy number data considerably increases the predictive performance of cognate gene expression relative to copy number alone (Supplementary Figure 2, Supplementary Table 3). We found that adding methylation driver genes led to an averaged R-squared increase of between 6% for LUSC up to 16% for BRCA when predicting cognate gene expression (Figure 3, Supplementary Figure 2).

**Figure 3:**
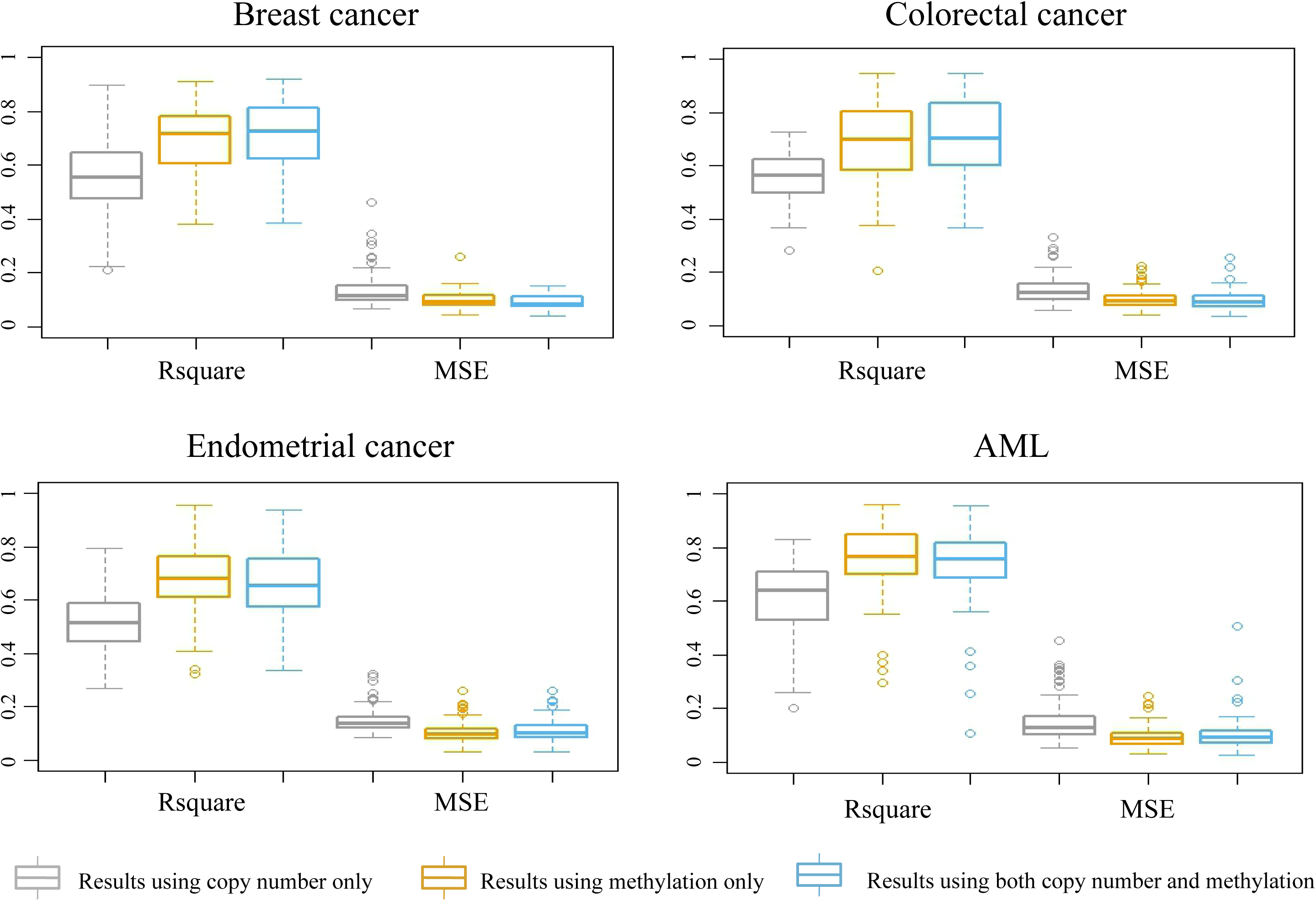
Boxplots representing the prediction performances of cancer driver genes predicting their target genes measured in two ways: R-square and mean squared error (MSE) obtained after running AMARETTO in three ways using only copy number data, only DNA methylation data and with both copy number and methylation data, shown for four cancer sites.

### Connecting AMARETTO modules reveals pancancer driver genes

To identify pancancer driver genes, we connected AMARETTO modules across 11 cancer sites in a network and identified significantly connected subnetworks. Our results show a module network with 2,693 edges between 713 modules (Supplementary Figure 3). Given this network, we detected 20 subnetworks containing between 9 and 74 modules (Figure 4, Supplementary Table 4). Among these subnetworks, seven represent all 11 cancer sites. The most heterogeneous one is Subnetwork 17, consisting of 11 modules each representing a different cancer. The least represented cancer site is LAML, with only 26 modules. It is also absent from 7 subnetworks reflecting most likely the difference between hematological and epithelial cancers. On the contrary, HNSC, LUSC, LUAD and BLCA, which are part of 19 subnetworks, are over-represented with more than 60 modules. Next, we focused on subnetworks that show high degrees of overlapping cancer driver genes in an effort to identify pancancer driver genes of important biological pathways. For example we identified a pancancer subnetwork enriched in cell cycle pathways (Figure 4, dark blue subnetwork, Supplementary note). Next, we describe two subnetworks that led to the identification of two novel pancancer driver genes: a subnetwork involved in smoking induced cancers and a subnetwork involved in ‘antiviral’ interferon-modulated innate immune response.

**Figure 4:**
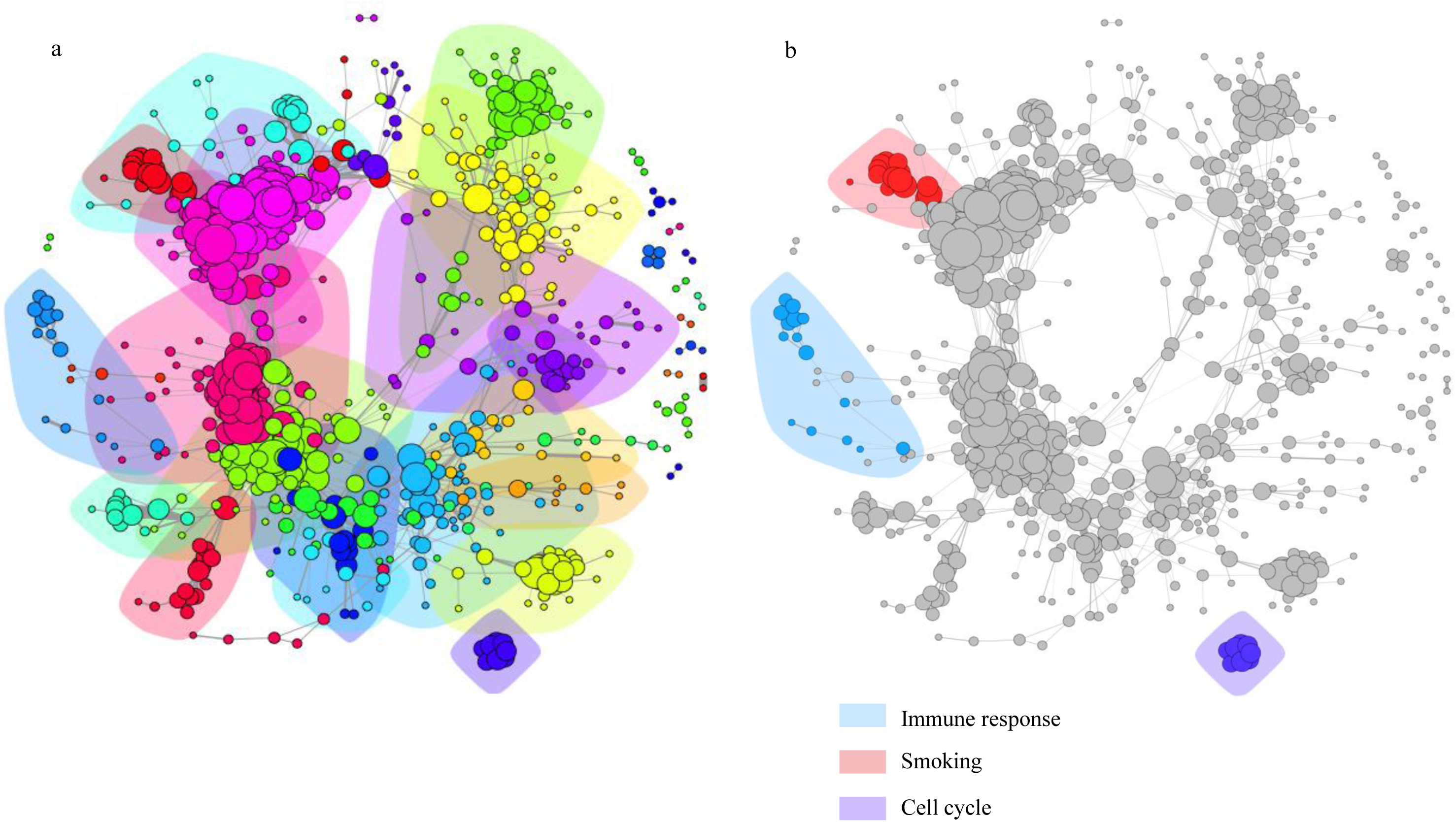
(a) Network representing all pancancer modules and 20 subnetworks. One node corresponds to one module from one cancer site. An edge between two modules represents a significant association between the module genes. The identified subnetworks are highlighted using different colors. (b) Three subnetworks are highlighted: the blue subnetwork is related to smoking, the red subnetwork captures immune response and the dark blue subnetwork captures the cell cycle.

### GPX2 is a driver of smoking induced cancers

We found a subnetwork containing 15 modules representing 8 different cancers that is significantly enriched in smoking induced pathways (Figure 4, blue subnetwork). Two smoking associated cancer sites, LUSC and HNSC, have three modules each in this subnetwork. This subnetwork contains one cancer driver gene GPX2, a glutathione peroxidase from the GPx family of genes, that is part of 8 modules across multiple cancer sites (Supplementary Table 5). GPX2 is hypo-methylated gene in all of the 11 cancers, except in HNSC where it is hyper or hypo-methylated in different patient subgroups. We found that several modules of the subnetwork are enriched in three smoking-related pathways (Supplementary Table 5). These particularly include genes involved in protection against chronic inflammation and asthma in lung cancers, as well as smoking-related gene expression signatures ^47-49^.

To verify the smoking association of this subnetwork, we used two gene signatures reflecting smoking damage, a xenobiotic metabolism signature, and an oxidative stress gene signature (Supplementary Table 6a). This analysis showed a significant correlation between GPX2 expression and xenobiotic metabolism for all cancers (p-value < 0.001, Figure 5a, Supplementary Figure 4, Supplementary Table 6a) and a significant correlation with oxidative stress for six cancer sites in this subnetwork (p-value < 0.001, Supplementary Figure 5, Supplementary Table 6a), suggesting a role of GPX2 in meditating cellular response to carcinogens in tobacco smoke ^50, 51^.

**Figure 5:**
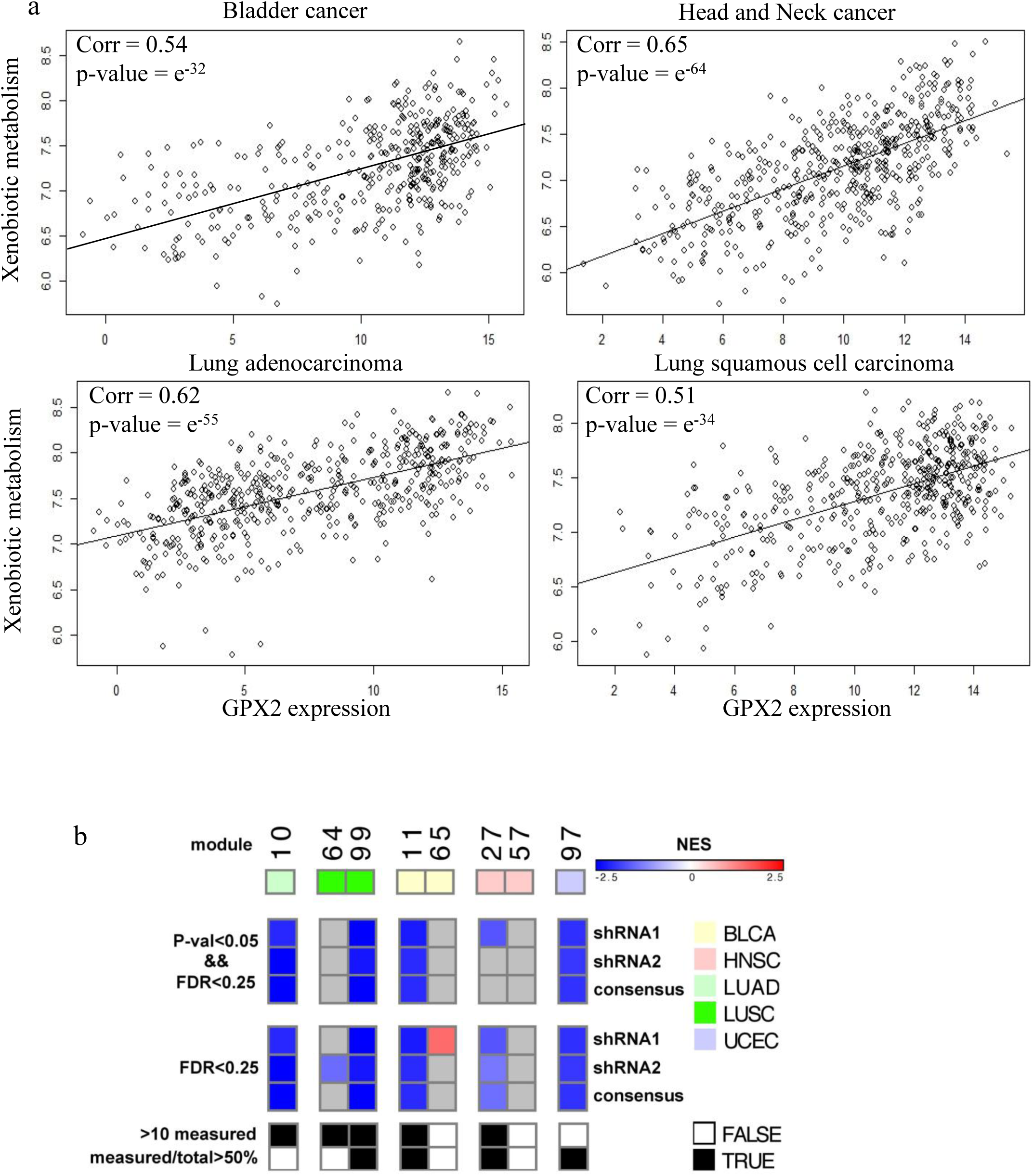
(a) Correlation between GPX2 expression and the averaged expression of a xenobiotic response gene signature showing significant correlation. (b) Heatmap showing the enrichments of the target genes of the 8 modules regulated by GPX2 derived from the 5 cancer sites (columns organized by cancer sites in following order: LUAD, LUSC, BLCA, HNSC, UCEC) in the three GPX2 knockdown experiments measured in the lung adenocarcinoma A549 cell line (rows: shRNA1, shRNA2, and consensus). Enrichments are represented by the GSEA normalized enrichment scores (NES). Only significant enrichments (top panel: p-value<0.05 and FDR<0.25; bottom panel: FDR<0.25) are shown (red: induced; blue: repressed; grey: not significant). The bottom panel shows the modules that contain at least 10 target genes or at least 50% of the target genes in LINCS.

Next, we correlated GPX2 expression with smoking data, available for BLCA, HNSC, LUAD and LUSC. Even in the presence of significant missing clinical data (Supplementary Table 6b), we found significant correlations with the number of smoked years (p-value = 0.011, corr=0.26) and pack years (p-value=0.003, corr=0.21) for HNSC (Supplementary Table 6b). We also found a significant association with GPX2 expression and smoking profile for HNSC (p-value < 0.001, Supplementary Figure 6), the most represented cancer within the subnetwork, and a borderline significant correlation for BLCA (p-value=0.05, Supplementary Figure 6).

To experimentally validate GPX2 as a driver of the smoking subnetwork, we interrogated the target genes of the 8 modules learned in the 5 cancer sites against publicly available genetic perturbation studies of GPX2 in the Library of Integrated Network-Based Cellular Signatures (LINCS) database ^28, 52^. We used Gene Set Enrichment Analysis (GSEA) to test for enrichment of the target genes of these modules in GPX2 knockdown experiments performed in the lung adenocarcinoma A549 cell line ^29, 53^. We observed that the target genes of these modules that are activated by induced GPX2 expression are significantly repressed upon knockdown of GPX2 in the A549 cell line. This expected behavior was observed in LUAD, and also consistent in the four other cancer sites, i.e., LUSC, BLCA, HNSC and UCEC (Figure 5, Supplementary Table 7), which confirmed our hypothesis that GPX2 is a causative driver of these smoking-related modules.

### OAS2 is a driver of ‘antiviral’ interferon-modulated innate immune response

We found a second intriguing pancancer subnetwork, Subnetwork 12, containing 15 modules representing 10 different cancers and enriched in interferon-inducible antiviral response pathways (Figure 4, red subnetwork). After ranking the 65 cancer driver genes in this subnetwork based on overlap, we identified one gene, OAS2, regulating 10 modules, and TRIM22 regulating 6 modules, 6 genes regulating 4 modules and 8 genes overlapping 4 modules (Supplementary Table 8). OAS2 is an interferon (IFN)-inducible enzyme that senses of double-stranded RNA (dsRNA) produced by viruses and subsequently activates RNAse L to destroy viruses^54^. Next, TRIM22 is also an interferon-inducible motif family antiviral protein, though its mechanism of viral repression is less clear^55, 56^. Interestingly, almost all of the cancer driver genes of this subnetwork present aberrant methylation patterns.

Using enrichment analysis, we found that the subnetwork is enriched by 21 different gene sets, most of which represented antiviral response and/or interferon inducible pathways (Supplementary Table 8). These included genes up or down-regulated in T-cells ^57^, dendritic cells ^58^, and other blood cell types ^59^. A strong enrichment was also found with an IFN gamma response gene set from the hallmark gene collection ^26^ and other IFN response gene sets were highly enriched ^60^. Using the IFNs database Interferome as a reference ^61^, we confirmed that pancancer driver genes of the immune subnetwork were related to response to IFNs, including all three IFN classes (Supplementary Figure 7). Despite its role in antiviral response, OAS2 was not differentially expressed between cancers harboring oncogenic viruses and those without detectable viruses, based on data for detection of viral transcripts in TCGA cancers provided ^62^ (data not shown).

Immune gene expression signatures in cancer can reflect the profile of mixed infiltrating tumor associated leukocyte (TAL) cell types within the tumor. To determine the TAL types associated with the IFN responsive signature, we inferred the levels of 22 TAL types in all TCGA cases using CIBERSORT ^30^, and tested the correlation of pancancer driver genes OAS2 and TRIM22 with each TAL type. Both OAS2 and TRIM22 were strongly correlated with M1 macrophage levels across all cancer types. This is consistent with the fact that these pro-inflammatory M1 tumor associated macrophages (TAMs) are activated by IFNγ in response to pathogen infection or cancer, and that M1 macrophage activation or ‘polarization’ coincides with upregulation of IFN-responsive genes ^63^. Interestingly, SP110, a pancancer driver gene of the IFN response subnetwork, is known to regulate macrophage gene expression and differentiation ^64, 65^.

We have recently reported that a molecular subtype of HNSC with high levels of M1 TAMs overexpresses CD274, the gene encoding PD-L1, a ligand for the CD8+ T cell-expressed immune checkpoint receptor PD-1 ^66^. This suggests that M1 TAM expression of PD-L1 may contribute to evasion of CD8+ T cell-mediated anti-cancer immunity, as previously reported ^63^. To investigate this further, we tested the correlation of OAS2, as a marker of the IFN-responsive/M1 TAM signature, with both ligands for the immune checkpoint receptor PD-1: CD274, the gene encoding PD-L1, and PDCD1LG2, encoding PD-L2. We observed a significant correlation for both genes and all cancer sites (p-values < 0.001, Supplementary Figure 8, Supplementary Table 9).

## Discussion

Here, we present a pancancer analysis using AMARETTO, an algorithm that addresses the challenge of integrating and interpreting multi-omics cancer data. AMARETTO first identifies cancer drivers, by considering genes with either DNA copy number or DNA methylation aberrations that have an effect on gene expression. AMARETTO then connects these cancer driver genes with target genes in the form of modules. Modules are subsequently associated with known biological pathways. We have shown that AMARETTO captures major biological pathways and reveals pancancer driver genes through network analysis.

AMARETTO focuses on DNA copy number and DNA methylation altered genes and their effect on gene expression, and does not model DNA somatic mutations for several reasons. First, somatic mutations do not necessarily affect gene expression, and additional data are required to be able to model the effect of a somatic mutation on expression, the key idea behind AMARETTO. Secondly, for each cancer site, besides the most significant mutated genes, many sequencing projects show that many genes are mutated in less than 5% of the cohort. These long tails create very sparse mutation data that do not add any predictive power in AMARETTO (data not shown). Future algorithmic work is needed to investigate how sparse somatic mutations can be integrated in a multi-omics framework like AMARETTO. Overall, AMARETTO is a complementary technique to identify cancer driver genes alongside methods focusing on distinguishing driver mutations from passenger mutations from DNA sequencing data such as MutSig ^67^.

Other computational methods have focused on identifying cancer driver genes using transcriptomics and multi-omics data. For example, ARACNE is a method that uses gene expression and a mutual information statistic to identify cancer driver genes through connecting transcription factors to their targets ^68, 69^. ARACNE is thus focused solely on gene expression data. CONEXIC is the most similar method to AMARETTO and uses a Bayesian strategy to connect cancer driver genes to their targets ^7^. We argue however that AMARETTO significantly improves upon CONEXIC. First, CONEXIC only takes into account DNA copy number changes, and does not model DNA methylation data. Our results show that DNA methylation driven genes are more predictive of transcription. Secondly, in a large benchmark, a previous comparison between AMARETTO and CONEXIC showed that in terms of R-square AMARETTO outperforms CONEXIC when predicting gene expression on unseen data ^70^. Thirdly, CONEXIC involves a large number of parameters and is more computationally demanding for large data sets ^70^.

In our results, we focused particularly on the biological implications of the two novel pancancer driver genes: GPX2 related to smoking and OAS2 related to IFN response. Regarding the former subnetwork, previous work has shown the importance of GPX2 in lung cancer. The GPX genes, glutathione peroxidases, are involved in protection of cells against oxidative stress and have been shown to be regulated by the Nrf2-pathway in lung. Activation of the Nrf2-pathway plays an important role in resistance to oxidant stress and its deregulation is one of the major causes of lung cancers ^71, 72^. Among the 5 GPX genes, expression of GPX2 has been shown to be related to smoking response, induced by Nrf2 activation in the lungs ^73^. GPX2 inhibits apoptosis in response to oxidative stress ^74^, such as may be caused by smoking. This subnetwork discovered by pancancer AMARETTO analysis indicates a key role for GPX2 in regulating gene expression in smoking-related cancers (LUAD, LUSC, HNSC, BLCA), and surprisingly, in UCEC, for which smoking is a protective factor. That GPX2 may block oxidative stress-induced apoptosis suggests that its inhibition may restore apoptosis, making it a potential drug target.

Next, the interferon response subnetwork showed that all of its predicted cancer driver genes represent genes that are expressed in response to IFNs. IFNs are cytokines that protect against cancer by activating innate immune inflammatory response to pathogens or cancer, triggering cancer cell death ^75^. We found that OAS2, overexpressed due to DNA hypo-methylation, was a major driver of IFN/immune related modules. While OAS2 is primarily implicated in mediating immune response to viruses ^76, 77^, most of the cancers overexpressing the IFN module do not have detectable oncogenic viruses ^62^. This is consistent with previous reports of an ‘antiviral’ gene expression profile marked by expression of OAS2 and other IFN responsive genes, in non-virus-related cancers ^78^. A similar ‘interferon-inducible antiviral response’ gene expression signature (Featuring OAS2, OAS1, IRF7, MX2 and ISG20, regulators of our IFN response subnetwork) was associated with response to expression of dsRNA derived from human endogenous retroviruses (HERVs) in ovarian and colorectal cancer cell lines, upon loss of DNA methylation-mediated repression of HERVs ^79, 80^. Given that OAS2 is a viral dsRNA sensor, and all of the IFN response subnetwork regulators were abnormally methylated, it is plausible that expression of this subnetwork reflects response to reactivation of HERVs, a frequent event in, and potential cause of, many cancers ^81^.

The IFN-response subnetwork was associated with levels of M1 TAMs across cancer types. TAMs include both pro-inflammatory M1 TAMs and anti-inflammatory M2 TAMs, both of which derive from M0 mature macrophages. M1 macrophage activation is stimulated by IFNγ, and is associated with expression of IFN-responsive genes such including OAS2 ^63, 82, 83^. Therefore, expression of IFN-response subnetwork genes may reflect infiltration of M1 TAMs, as part of innate response to cancer. Such inflammatory responses are generally considered to be anti-tumorigenic, indeed, stimulation of inflammatory response by treatment with IFNα is used therapeutically to stimulate anti-cancer immune response ^63, 84^. The current paradigm asserts that M2 TAMs are immunosuppressive and their levels are generally associated with poor prognosis in cancer, as they promote invasion, metastasis and therapy resistance. Conversely M1 TAMs that kill tumor cells are associated with prolonged survival ^85-87^. On the other hand, tumors can modulate TAMs to express pro-oncogenic factors. We observed that OAS2 expression is correlated with expression of CD274, the gene encoding PD-L1, a ligand for the CD8+ T cell-expressed immune checkpoint receptor PD-1 that suppresses CD8+ T cell-mediated anti-cancer immunity. Indeed, IFN induces expression of PD-L1 during chronic inflammation or viral infections, dampening CD8+ T cell response ^88, 89^. PD-L1 is expressed by TAMs as well as tumor cells, and emerging evidence indicates that tumors modulate TAMs to express high levels of PD-L1 ^63, 90^. Our findings indicate that PD-L1 expression is particularly correlated with M1 TAMs, as opposed to M0 or M2 TAMs, indicating a novel immunosuppressive role of M1 TAMs.

Monoclonal antibodies targeting PD-L1 or PD-1 can restore CD8+ T cell cytotoxic anti-cancer immunity, suggesting a potential therapeutic opportunity for patients displaying the IFN signature. Indeed, an IFN signature has recently been shown to be favorably predictive of response to PD-1 blockade ^91, 92^ and recent experimental evidence indicates that IFN signaling, particularly IFN gamma signaling upregulates PD-L1 and PD-L2 expression in cancer, both in vitro and in vivo ^93^. A cancer driver gene such as OAS2 or TRIM22 may help to distinguish between patients that will benefit for PD-1/PD-L1 immunotherapy, and those for whom ineffective treatment may cause autoimmune side effects ^94^. AMARETTO has hereby enabled us to identify OAS2 and TRIM22 as epigenetically deregulated cancer driver genes within a pan-cancer IFN responsive pathway that provides novel biological insight into tumor-immune interactions that may have implications for immunotherapy.

In summary, pancancer AMARETTO allows identifying cancer driver genes for major hallmark cancer pathways, a pancancer driver gene involved in smoking induced cancers and a pancancer driver gene of immune response. AMARETTO thus provides a computational method for cancer driver gene identification in a multi-omics setting and might lead to novel drug targets.

## Acknowledgements

We thank The Cancer Genome Atlas (TCGA) for making the data publicly available.

## Author contributions

OG conceived and designed the study. OG, KB and MC developed the algorithm, performed experiments and analyzed the data. KB and AG analyzed CIBERSORT data and interpreted results. TC and NP performed experiments and carried out analysis using LINCS data. All authors contributed in manuscript drafting and revision.

## Supplementary Figures

Supplementary Figure 1: Number of regulator genes (genes that regulate at least one module from one cancer type) identified in AMARETTO by only copy number, only DNA methylation or both.

Supplementary Figure 2: Boxplots representing the prediction performances (mean squared error MSE and R-square) obtained after running AMARETTO using copy number data only, methylation only or both copy number and methylation on the 11 cancer sites.

Supplementary Figure 3: Overview of the module network (upper figure), the pancancer module network (on the bottom, left) and a zoomed view of intriguing subnetworks. The nodes of the graph are all the modules across all cancers sites. Their size depends on the node degree (number of incident edges). An edge between two modules stands for a significant association between them (measured through the minus log-transformation of the adjusted p-value, which also defines the edge thickness). For the top figure, the node color depends on the associated cancer site, for the two bottom figures, the node color depends on the subnetwork it belongs to.

Supplementary Figure 4: Correlations between GPX2 expression and the averaged expression of an oxidative response gene signature for all cancer sites.

Supplementary Figure 5: Correlations between GPX2 expression and the averaged expression of a xenobiotic response gene signature for all cancer sites.

Supplementary Figure 6: Boxplots representing the association between GPX2 expression and smoking profile for BLCA and HNSC cancers.

Supplementary Figure 7: Venn diagram representing the number of genes regulating the immune response subnetwork and induced by IFNs of type I, II or III.

Supplementary Figure 8: Scatterplots representing the associations between OAS2 expression and PD-L1 expression on the left, and PD-L2 expression, on the right for all cancer sites.

## Supplementary Tables

Supplementary Table 1: (a) The top 50 most selected driver genes regulating modules enriched in major pathways of cancer including angiogenesis, hypoxia, EMT, cell cycle, immune response, apoptosis, metastases, integrin signaling and EGFR across all cancer sites, (b) genetic and epigenetic alterations of the top drivers across all cancer sites and (c) comparison of average number of enriched gene sets per module in 100 random permutations vs. the actual modules per cancer.

Supplementary Table 2: Number of modules regulated by driver genes for all cancer types. Only the top regulator genes, ranked according to the total number of regulated modules across all cancer sites (last column) are represented.

Supplementary Table 3: Prediction performances (R-square and mean squared error MSE) obtained after running AMARETTO using copy number data only, methylation only or both copy number and methylation on the 11 cancer sites. The two last columns indicate the R-square and MSE increase when adding methylation data.

Supplementary Table 4: Distribution of the modules per cancer (column) and subnetwork (row). The last column indicates the total number of modules within a subnetwork.

Supplementary Table 5: Cancer driver genes and enrichment results of the smoking subnetwork ranked by number of modules each driver gene is participating in.

Supplementary Table 6: (a) Oxidative and xenobiotic gene signatures provided by the GO ontology and used to measure the association of GPX2 expression with smoking. (b) Correlations and p-values measuring the association between smoking related data (smoking profile, number of smoked years and pack years) and GPX2 expression across all cancer sites with enough data (BLCA, HNSC, LUAD, LUSC). The percentage of missing data is also indicated.

Supplementary Table 7: Experimental validation of the modules and their target genes regulated by GPX2 as a causative driver of the pancancer smoking subnetwork. Table shows overall GSEA enrichment results of the three perturbation experiments upon GPX2 knockdown in the lung adenocarcinoma A549 cell line derived from LINCS (columns: consensus, shRNA1, shRNA2) in each of the 8 modules regulated by GPX2 in the 5 cancer sites (rows: modules organized by the 5 sites in the following order: LUAD, LUSC, BLCA, HNSC, UCEC). On top are shown the significance levels (p-values and FDR values; green if p-value<0.05 and FDR<0.25, yellow if FDR<0.25) and below, the normalized enrichment scores (NES; blue if repressed, red if induced). On the right are the sizes of the signatures, from left to right: number of genes in the modules, number of genes that are part of the LINCS landmark and bing genes, and the numbers of those genes that are identified as ‘leading edge’ genes driving the GSEA enrichment scores in the three experiments (consensus, shRNA1, shRNA2).

Supplementary Table 8: Cancer driver genes and enrichment results of the immune response subnetwork ranked by number of modules each driver gene is participating in.

Supplementary Table 9: Correlations and p-values for Pearson test measuring the association between OAS2 expression and PD-L1/PD-L2 expression for all cancer sites.

Supplementary Table 10: Cancer driver genes and enrichment results of the histone subnetwork ranked by number of modules each driver gene is participating in.

